# Neuroanatomy of the clitoris

**DOI:** 10.64898/2026.03.18.712572

**Authors:** Ju Young Lee, Demi Alblas, Adam Szmul, Daniël Docter, Hector Dejea, Yousif Dawood, Jermo Hanemaaijer-van der Veer, Alexandre Bellier, Theresa Urban, Joseph Brunet, David Stansby, Joanna Purzycka, Ruikang Xue, Claire L Walsh, Peter D Lee, Paul Tafforeau, Roelof-Jan Oostra, Robert CJ Kanhai, Joseph Jacob, Joris AM van der Post, Otto Bleker, Stephanie Both, Judith AF Huirne, Bernadette S de Bakker

**Author notes:** Corresponding author (J.Y.L), (B.S.d.B.).

## Abstract

The clitoris is one of the least studied organs of the human body. The detailed anatomy of the clitoris is challenging to address through a gross dissection, as most of its parts are embedded internally, surrounded by pubic bone and several pelvic organs. While clinical imaging methods such as magnetic resonance imaging can capture the gross 3D morphology, they lack the spatial resolution required to resolve the detailed structures. In this study, we generated micron-scale computed tomography images of the female pelvises, leveraging a synchrotron radiation X-ray source. This unique data revealed the complex trajectory of the dorsal nerve of the clitoris, the main sensory nerve of the clitoris. Notably, the nerve trunks within the clitoral glans were revealed, with the maximum diameter ranging from 0.2 to 0.7 mm. They showed a tree-like branching pattern projecting towards the surface of the glans. We also revealed that some branches of the dorsal nerve of the clitoris ramify to innervate the clitoral hood and mons pubis. Finally, the posterior labial nerve, a branch of the perineal nerves, was shown to innervate the surroundings of the clitoris and the labial structures. These findings have an immediate impact on operations performed around the vulva area, such as gender-affirmation surgery and reconstruction surgery after genital mutilation.

## Introduction

The clitoris is a unique organ in the female body, responsible for sexual pleasure. Early references to the clitoris can be traced back to ancient times, exemplified by the writings of Aristotle [1, 2]. However, the cultural taboo around female sexuality has hindered its scientific investigation for centuries [3]. In 16^th^ century, the clitoris was described as the “shameful member (membre honteux)” by a French anatomist [4]. The standard anatomy textbook did not include the clitoris until the 20th century. When it was introduced in the 38th edition of Gray’s Anatomy, it was erroneously referred to as ‘a small version of the penis’ [5].

Seminal works from O’Connell [6-8] described a comprehensive anatomy of the clitoris. Using magnetic resonance imaging, it was possible to appreciate the internal structure of the female genitalia in 3D, including the erectile tissues. The erectile tissue of the female genitalia consists of corpus cavernosa and corpus spongiosum, similar to male genitalia. Instead of surrounding the urethra like the corpus spongiosum in a penis, the female corpus spongiosum is split into two bulbs and flanks the vaginal wall laterally. These works also revealed that the size of the clitoris is at least twofold bigger than depicted in anatomical textbooks [9, 10]. The clinical relevance was high, as it provided clinicians insights to better preserve the clitoris during pelvic operations.

Recent studies used methods ranging from gross dissection and microscopic imaging to investigate clitoral nerves [11]. Gross dissections on cadavers demonstrated the primary sensory pathway, the dorsal nerve of the clitoris (DNC), which emerges from the perineal membrane bilaterally [12, 13]. Histological studies revealed that the innervation density of the clitoris is several times greater than that of the penis, with reports ranging from 6- to 15-fold [14-16]. However, the full 3D trajectory from the clitoral nerve is yet to be described, as the nerves are largely embedded deep inside the body and are therefore more challenging to dissect compared to the penis.

To resolve the complex neuroanatomy of the clitoris, a non-invasive imaging method with micrometre-scale resolution capability is needed. In this research, we used Hierarchical Phase-Contrast Tomography (HiP-CT), a synchrotron-based X-ray phase-contrast microtomography method [17, 18]. A synchrotron is a cyclic particle accelerator that generates high-energy, coherent X-rays. Synchrotron light sources can be used to scan 3D images with voxel sizes down to the micrometre scale. The coherent X-rays enable the phase-contrast technique, providing sufficient contrast to determine biological compartments from soft tissue.

We scanned two postmortem female pelvic samples. The 3D trajectory of the nerves innervating the clitoris is demonstrated in unprecedented detail. Our results characterise the complex tree-like branching pattern of the DNC within the clitoral glans and identify its additional innervation to the clitoral hood and mons pubis. Furthermore, we provide a 3D trajectory of the posterior labial nerves (PLN), projecting to the clitoral hood and the labia. These findings will help clinicians when performing operations around the vulva area, such as child delivery, gender-affirmation surgery, and reconstruction surgery after genital mutilation.

## Results

### 1) Dorsal nerve of the clitoris at the clitoral body

Figure 1 demonstrates the main trajectory of the DNC, the primary sensory nerve of the clitoris. The DNC is a branch of the pudendal nerve, which innervates the pelvic region. It follows the shape of the corpus cavernosum, from the crura, the two-legged structure, to the clitoral body, where the two crura merge (Figure 1A). The nerves from the left crus stay at the 1- to 2-o’clock positions, and the nerves from the right crus stay at the 10- to 11-o’clock position (Figure B-D), in alignment with previous studies [12, 13, 19, 20]. The DNC extends to the glans of the clitoris, which is the external part of the clitoris densely populated with sensory receptors [16].

**Figure 1.**
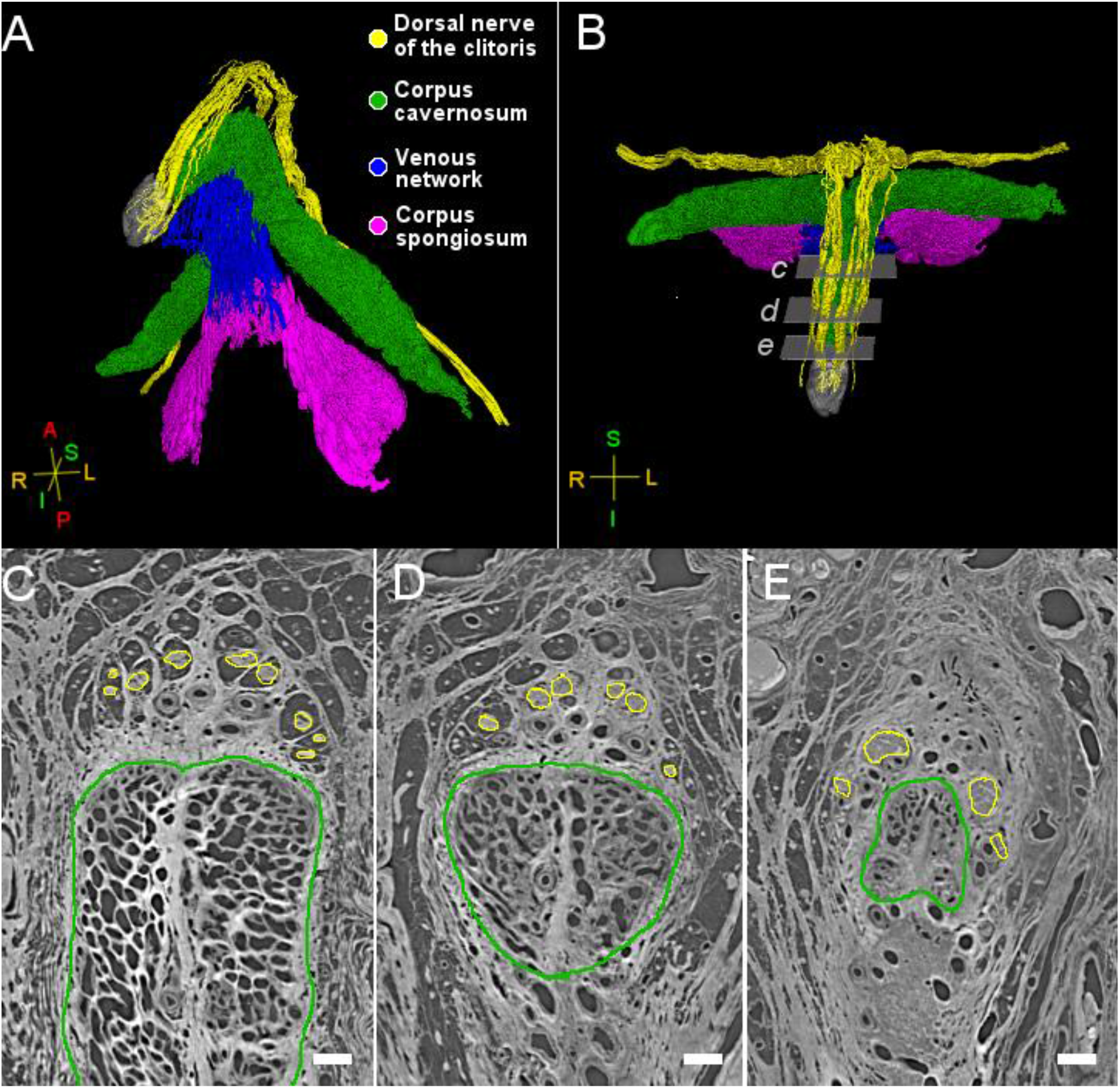
Dorsal nerve of the clitoris (DNC). (A, B). Overview of the clitoris. The colour coding is as follows: the dorsal nerve of the clitoris in yellow, the corpus cavernosum in green, the venous network in blue, the corpus spongiosum in magenta and the glans in transparent grey. Axes are on the left corners, with L, R, S, I, A, and P indicating left, right, superior, inferior, anterior, and posterior directions. (C-E). Cross sections of the clitoral body from proximal to distal. Yellow outlines indicate the DNC bundles. The cross sections are taken from the planes indicated in panel B (c-e). Scale bar = 1 mm. The width of the planes in Panel B is 1 cm.

### 2) Dorsal nerve of the clitoris at the clitoral glans

The DNC extends beyond the distal end of the corpus cavernosum into the clitoral glans. While traditional gross dissection is typically limited to the proximal boundary of the clitoral glans, HiP-CT images revealed the internal nerve trajectories of the clitoral glans with extensive branching projecting to the glans surface. Five nerve trunks were identified in both subjects (Figure 2, Supplementary Figure 1). The nerve trunks projected from medial towards the lateral surface of the glans (Supplementary Video 1), similar to the branching pattern in the penile glans [21]. The nerve trunks from left and right did not cross the midline, maintaining a lateral trajectory to innervate each hemisphere of the clitoral glans. The maximum diameter from each nerve trunk ranged from 230 µm to 700 µm (mean = 423.46 µm, SD = 159.64 µm, Supplementary Table 1).

**Figure 2.**
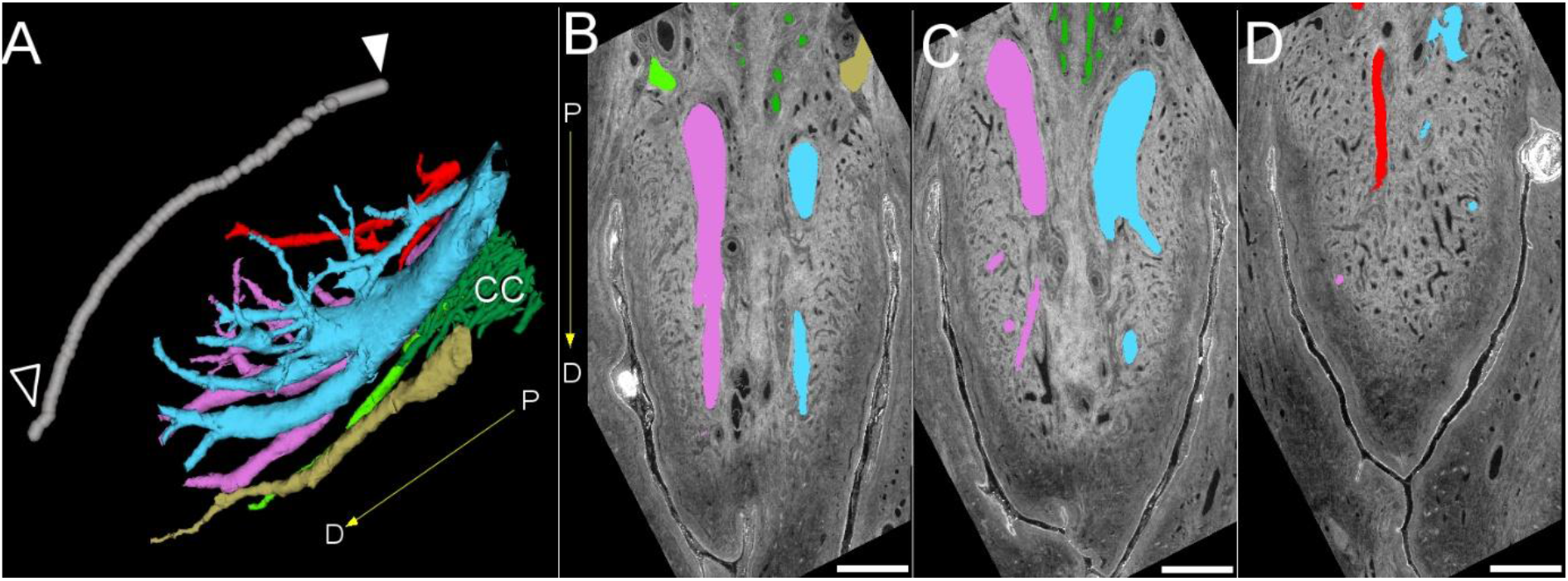
Dorsal nerve of the clitoris within the glans of the clitoris. A. Five large nerve trunks are shown within the glans, labelled in different colours. The axis which indicates the direction for distal (D) and proximal (P) ends of the glans is 3 mm long. The grey line indicates the contour of the glans surface. The filled arrowhead is on the proximal start of the glans, and the open arrowhead is on the distal end leading to the frenulum. The corpus cavernosum is labelled ‘CC’. (B-D). Sections along the distal and proximal ends of the glans are shown with an overlay of five nerve trunks with the same colour coding as panel A. Scale bar = 1 mm.

### 3) Dorsal nerve of the clitoris in the clitoral hood

In addition to clitoral glans innervation, we identified DNC branches that ramify superiorly at the angle of the corpus cavernosum (Figure 3). These branches travel towards the pubic symphysis and turn back ventrally, making an inverted U-shaped trajectory, seen from the sagittal view. This is in line with prior studies that described DNC within the suspensory ligament, a connective tissue between the clitoral body and the pubic symphysis [12, 22]. The branches terminate in different locations within the mons pubis and the clitoral hood.

**Figure 3.**
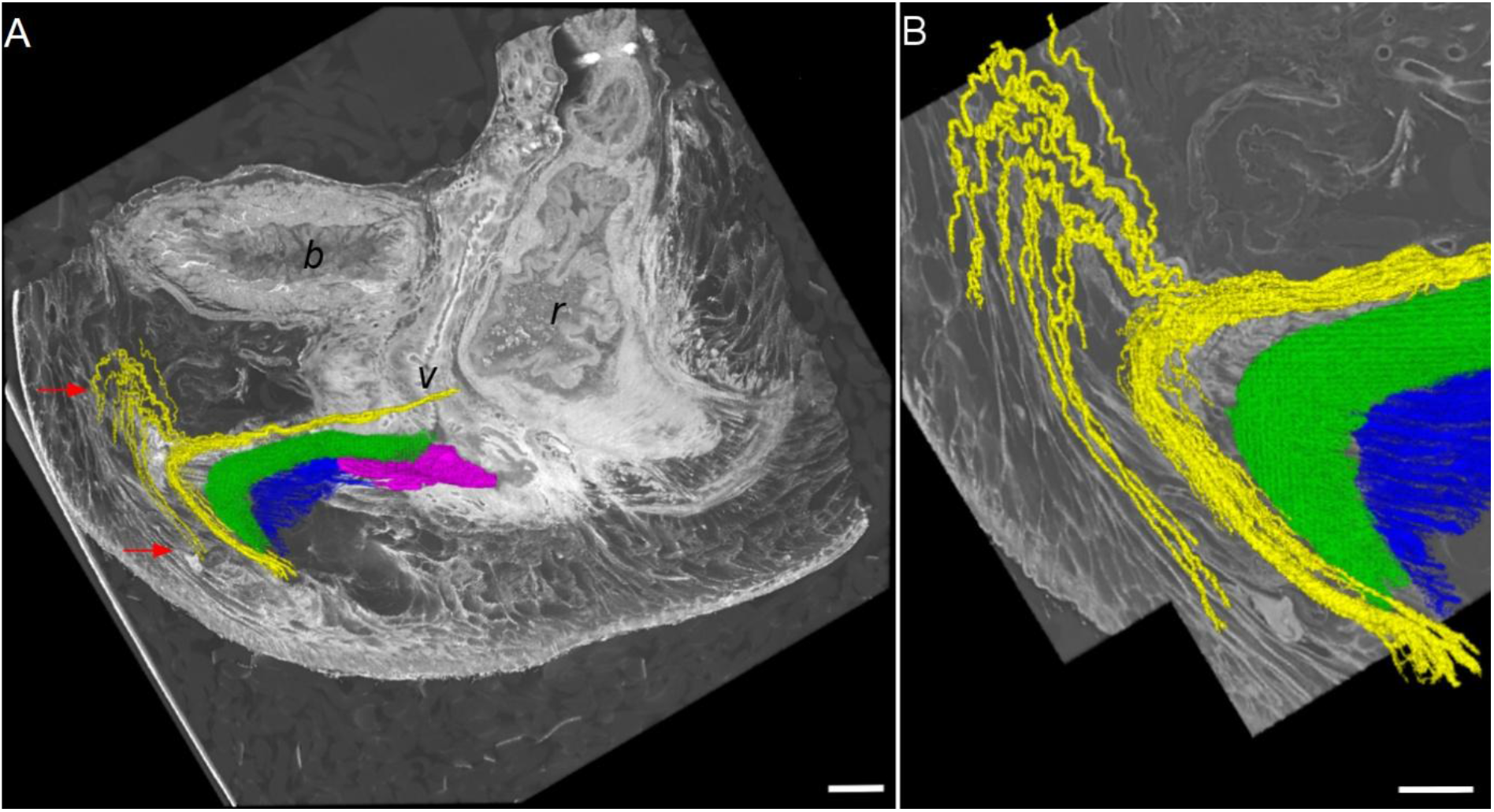
The dorsal nerve of the clitoris also innervates the mons pubis and the clitoral hood. A. Sagittal view showing the trajectory of the dorsal nerve of the clitoris. Red arrows indicate the innervation of the mons pubis (top) and the clitoral hood (bottom). The bladder, vagina and rectum are labelled as ‘b’, ‘v’ and ‘r’, respectively. The colour coding is as follows: the dorsal nerve of the clitoris in yellow, the corpus cavernosum in green, the venous network in blue, and the corpus spongiosum in magenta. Scale bar = 1 cm. B. Zoomed-in view of panel A. Scale bar = 5 mm.

### 4) The posterior labial nerve innervates labia and the surroundings of the clitoral body

Figure 4 demonstrates the innervation of the posterior labial nerve (PLN), a branch of the perineal nerve. The PLN travels inferior to the corpus spongiosum and projects towards several regions. While the PLN is known to innervate the labia majora and minora [11], we found that the PLN also provides innervation to the lateral aspects of the clitoral body.

**Figure 4.**
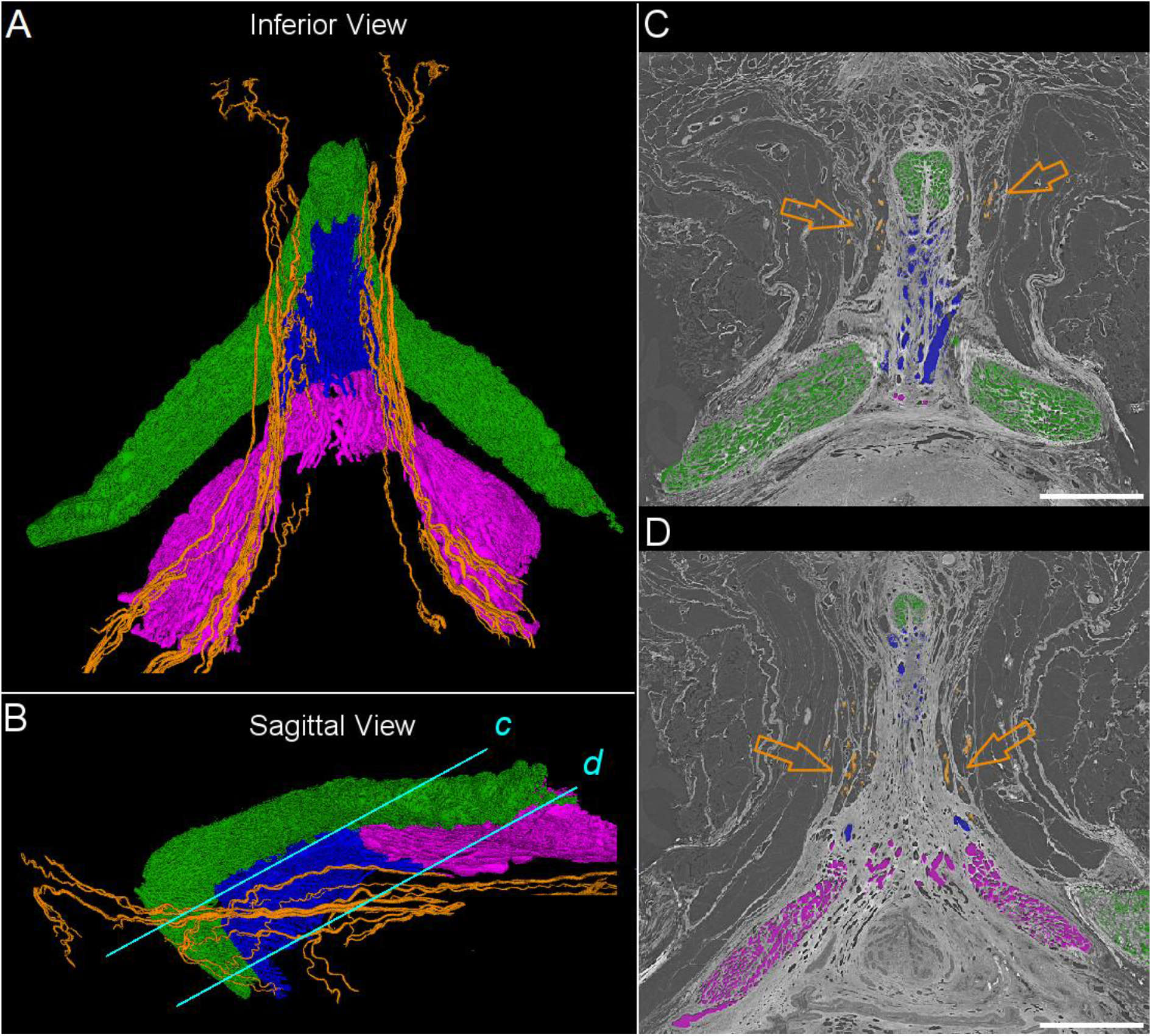
The posterior labial nerve (PLN) innervates the surrounding region of the clitoris. (A-B). An inferior (A) and sagittal (B) view demonstrating the trajectory of the PLN shown in orange, the corpus cavernosum in green, the venous network in blue, and the corpus spongiosum in purple. (C-D). Cross sections from lines indicated in panel B. The PLN branches are annotated with arrows. The colour coding is identical to panels A and B. Scale bars = 1 cm.

## Discussion

This study presents a comprehensive 3D map of the clitoral nerve structures, leveraging state-of-the-art X-ray imaging techniques. From high-resolution images of post-mortem human pelvises, we delineated a complete trajectory of the DNC, the main sensory nerve of the clitoris. In particular, 3D visualisation of the nerve trunks within the clitoral glans is revealed. Furthermore, we revealed that some branches of the DNC innervate the mons pubis and the clitoral hood. We also showed that the PLN, branches of the perineal nerve, not only project to the labia but also to the surroundings of the clitoral body.

This study addresses the knowledge gap in the neuroanatomy of the female genitalia. While the innervation of the penile glans has been documented for several decades [21], the clitoral glans remains comparatively understudied compared to the penile glans in the scientific literature between the years 2002 and 2022 [3]. This disparity is partly due to the technical challenge of investigating the clitoral glans for its smaller size.

Prior research described that the DNC ‘gradually diminishes’ as it approaches the glans [12]. However, this observation is likely due to the difficulty of following the DNC further into the glans using gross dissection. As a result, medical illustrations often depict the clitoral glans as having a sparse nerve supply [23]. By utilising 3D imaging with micrometre-scale voxel sizes, we have demonstrated that the DNC does not diminish but instead exhibits a complex tree-like branching pattern within the clitoral glans.

This study is highly relevant for the management of female genital mutilation (FGM) survivors, which affects over 230 million women globally [24]. The most widely practised FGM procedures are type 1, which removes the glans and the clitoral hood, and type 2, which removes the glans and labia minora. A comprehensive understanding of the neural pathways of the clitoral glans is essential for clinicians performing reconstructive surgeries. Longitudinal data indicate that approximately 22% of women who undergo clitoral reconstruction experience a post-operative decline in orgasmic experience [25, 26]. Our study provides the anatomical foundation necessary to investigate the physiological mechanism behind these sensory changes and potentially refine the surgical technique [27].

Our findings also suggest a need to revisit the definition of the “danger zone” in female genital cosmetic surgeries. In recent years, popularity for such surgeries has dramatically increased, exemplified by a 70% increase in labiaplasty from 2015 to 2020 [28]. Labiaplasty is a procedure to remodel the labia, which often involves incisions in the clitoral hood [29]. The “danger zone” has been suggested to help surgeons avoid nerve damage during the operation. While current literature includes the clitoral hood as a risk area [30, 31], our study shows some branches of the DNC extending beyond the defined danger zone, branching out to innervate the mons pubis. Recognition of these distal nerve branches may assist surgeons in refining operative techniques.

Several limitations of the present study must be acknowledged. First, the sample size was limited, which may not fully account for the spectrum of anatomical variability present in the general population. Second, the specimens utilised were obtained from postmenopausal donors; consequently, the findings may not fully reflect the morphology of premenopausal individuals. Finally, this work focused specifically on somatic sensory innervation and did not include a mapping of the autonomic nervous system, such as the cavernous or spongious nerves.

Future research incorporating larger, age-diverse cohorts is necessary to provide an exhaustive representation of clitoral innervation. Furthermore, studies should address the functional properties of the neurons of the clitoris using methods such as immunohistochemistry to identify both sensory and autonomic fibres. Recent studies on mice found that the clitoral glans has 16 times the density of mechanosensory neurones as the penile glans [16]. In human studies, a higher myelination ratio has been observed in the clitoris compared to the penis [15]. Future investigation using molecular markers will be crucial in determining the specific functionality of the neurons we have visualised in the clitoral glans, clitoral hood, labia and beyond [32].

In summary, this study provides a 3D neuroanatomical map of the clitoris, using high-resolution imaging to overcome previous methodological limitations. By tracing the complete trajectory of the DNC into the clitoral glans and identifying extensive innervation within the clitoral hood and mons pubis, these findings resolve a long-standing information gap in scientific literature. The detailed mapping of these neural pathways offers a critical framework for clinicians, particularly in refining surgical approaches for FGM reconstruction and genital cosmetic procedures. Ultimately, this work establishes an important anatomical foundation that facilitates future investigations into the functional and molecular characteristics of clitoris innervation.

## Methods

### Sample preparation

Two post-mortem pelvic samples (ages = 59, 69) were acquired through the body donor program of the Amsterdam University Medical Center. The donation of the bodies, from which the samples were extracted, adhered to Dutch legislation and the regulations set forth by the medical committee of the Amsterdam UMC (METC-number 2024.0486). The bodies were preserved with a 10% formalin fluid. To isolate the pelvic region, transections were made just superior to the iliac crest and at the level of the proximal one-third of the femur. To preserve the entire clitoris and as much of the surrounding anatomy as possible, a large sample was carefully dissected free from the bones. Anteriorly, the sample consisted of all the soft tissue within the bony pelvic triangle defined by the pubic symphysis superiorly and the inferior pubic ramus laterally. Posteriorly, the sample extended to an imaginary coronal plane through the midline of the rectum. Laterally, all excess tissue was removed. Superiorly, the sample followed an imaginary transverse plane through the uterus and bladder.

The tissue embedding protocol is explained extensively in previous papers [17, 18]. Briefly, the dissected samples were dehydrated in increasing ethanol concentration baths of 50%, 60% and, finally, 70% ethanol and mounted in a standardised container, using agar gel. To avoid artefacts from air bubbles, the samples were put in the vacuum chamber at -.09 MPa for 10 minutes several times (minimum 4 times). Between the vacuum steps, air bubbles were manually removed. In addition, two rounds of in-line degassing were performed.

### Image acquisition

HiP-CT was performed at the BM18 beamline of the European Synchrotron Radiation Facility following established protocols [33]. Imaging was carried out using a polychromatic X-ray beam, filtered using sapphire and silver, resulting in an average photon energy of approximately 100 keV.

Samples were scanned at two voxel size settings: 20 µm and 2 µm (Supplementary Table 2). The field of view (FOV) of the 20 µm voxel size covered the entire specimen, and the FOV of the 2 µm voxel size covered the clitoral glans (Supplementary Figure 2). Detection was performed using a gadolinium aluminium gallium garnet scintillator with reflective coating, coupled to an IRIS 15 camera (Teledyne Photometrics, United States).

### Image reconstruction

3D reconstruction was performed using Nabu, an open-source software for tomographic reconstruction, in combination with the night_rail framework [34, 35]. The reconstruction was executed on the ESRF high-performance computing cluster using a fully automated workflow. Briefly, the projection data were first corrected for dark-field and flat-field variations using images from the reference container filled with 70% ethanol. Single-distance phase retrieval was then applied using the Paganin algorithm, combined with an unsharp mask filter [36]. Tomographic reconstruction was performed using filtered back-projection. Post-reconstruction processing included 16-bit reconstructed volumes ranging from 38 GB to 1730 GB.

### Image segmentation

Nerves were segmented using the following approaches.

For the 20 µm voxel-size images, two machine learning algorithms were employed. First, we used a pre-trained no-new-U-net (nnU-Net) [37]. The model was trained on synchrotron data of the human lung acquired from the same beamline using the same protocol [38]. Four additional iterations of annotation and training were used to adapt the nnU-Net for the clitoris dataset. About 50 hours of manual annotation were needed for additional annotation. As a post-processing step, isolated island components were filtered out. The result showed the large nerves of DNC (Figure. 1A).

In addition, small branches of nerves of the DNC and PLN (Fig. 3, Fig. 4) were segmented using the Segment Anything Model [39]. Visually identified incomplete nerves were extended using the quick select and interpolate tools from the Webknossos platform [40].

From the 2 µm voxel-size images, the nerve trunks of the glans were segmented (Fig. 2). The data was downsampled by a factor of four. The downsampling did not result in the loss of nerve branches, as branches with thickness below 70 µm were difficult to visibly trace in the original scale. Similar to the above, the Segment Anything Model [39] was used within the Webknossos platform.

The corpus cavernosum, venous network, and corpus spongiosum were segmented from 20 µm voxel images using the following approach. First, the ROI was downsampled by a factor of 4. Then, a rough 3D binary mask was manually generated using the 3D brush tool using open-source software, ITK-SNAP [41]. The binary mask was applied to select each structure, followed by thresholding.

### Quantitative analysis

Quantitative analysis was conducted on the nerve trunks within the glans. Using the maximum inscribed ball with the MorphlibJ plugin [42] within Fiji software [43], the maximum diameter was calculated for each nerve trunk. The average and the standard deviation were calculated from each subject.

## Data Availability

All reconstructed volumes are openly accessible through the Human Organ Atlas repository (https://human-organ-atlas.fr).

## Acknowledgement

We thank Maeva Badre, Celine Brockmann and Elcin Tunçkol for feedback on the manuscript. This research was supported by the Amsterdam University Medical Centre, the European Synchrotron Radiation Facility (BM18 Beamtimes MD1290 and MD1389), University College London, and the Chan Zuckerberg Initiative (Grants 2022-316777 and 2024-009938).

## Author contributions

Contribution according to CRediT system: **J.Y.L**.: Investigation, Data curation, Methodology, Software, Formal analysis, Writing—original draft, **D. A**.: Conceptualization, Investigation, **A.S**.: Software, **D.D**.: Methodology, **H.D**.: Data curation, Methodology, **Y.D**.: Conceptualization, **J.H.V**: Project administration, **A.B**.: Conceptualization, Validation, **T.U**.: Data curation, Methodology, **J.B**.: Data curation, Methodology, **D.S**.:Software, Data curation, **J.P**.: Methodology, Project administration, **R.X**.: Project administration, **C.L.W**.: Methodology, Project administration, Funding acquisition, **P.D.L**.: Funding acquisition, Resource, **P.T**.: Methodology, Funding acquisition, Resources, **R.J.O**.:, **R.C.J.K**.: **J.A.P**.:, **O.B**.:, **S.B**.: Conceptualization, Supervision, **J.A.H**.: Resource, Supervision, **B.S.d.B**.: Resource, Project administration, Supervision

Everyone was involved in Writing – review & editing

The dissections were collectively performed by A.W. Kastelein, Y. Dawood and D. Alblas.

