## Supplemental Information for "Neuroanatomy of the clitoris"

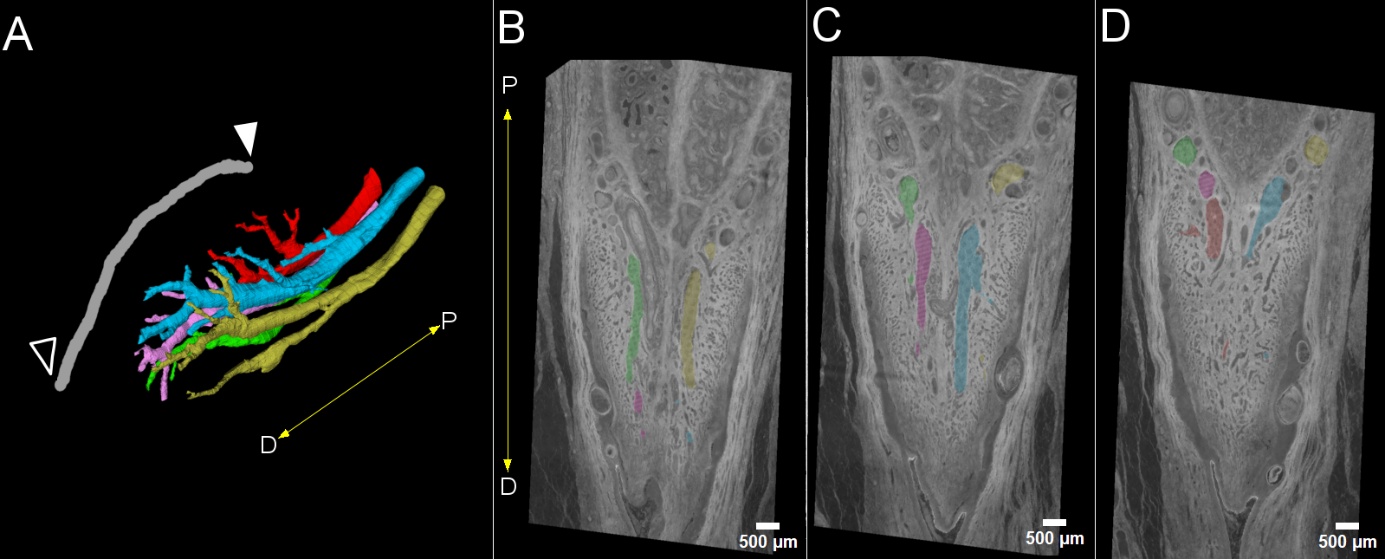

Supplementary Figure 1. Dorsal nerve of the clitoris within the glans of the clitoris from subject 2. A. Five large nerve bundles are shown within the glans, labelled in different colours. The axes indicate the direction for distal (D) and proximal (P) ends. The filled arrowhead indicates the proximal start of the glans. The open arrowhead indicates the distal end leading to the frenulum. (B-D). Sections along the distal and proximal ends of the glans are shown with an overlay of five nerve trunks, each in different colours. Scale bar = 0.5 mm

Supplementary Figure 2

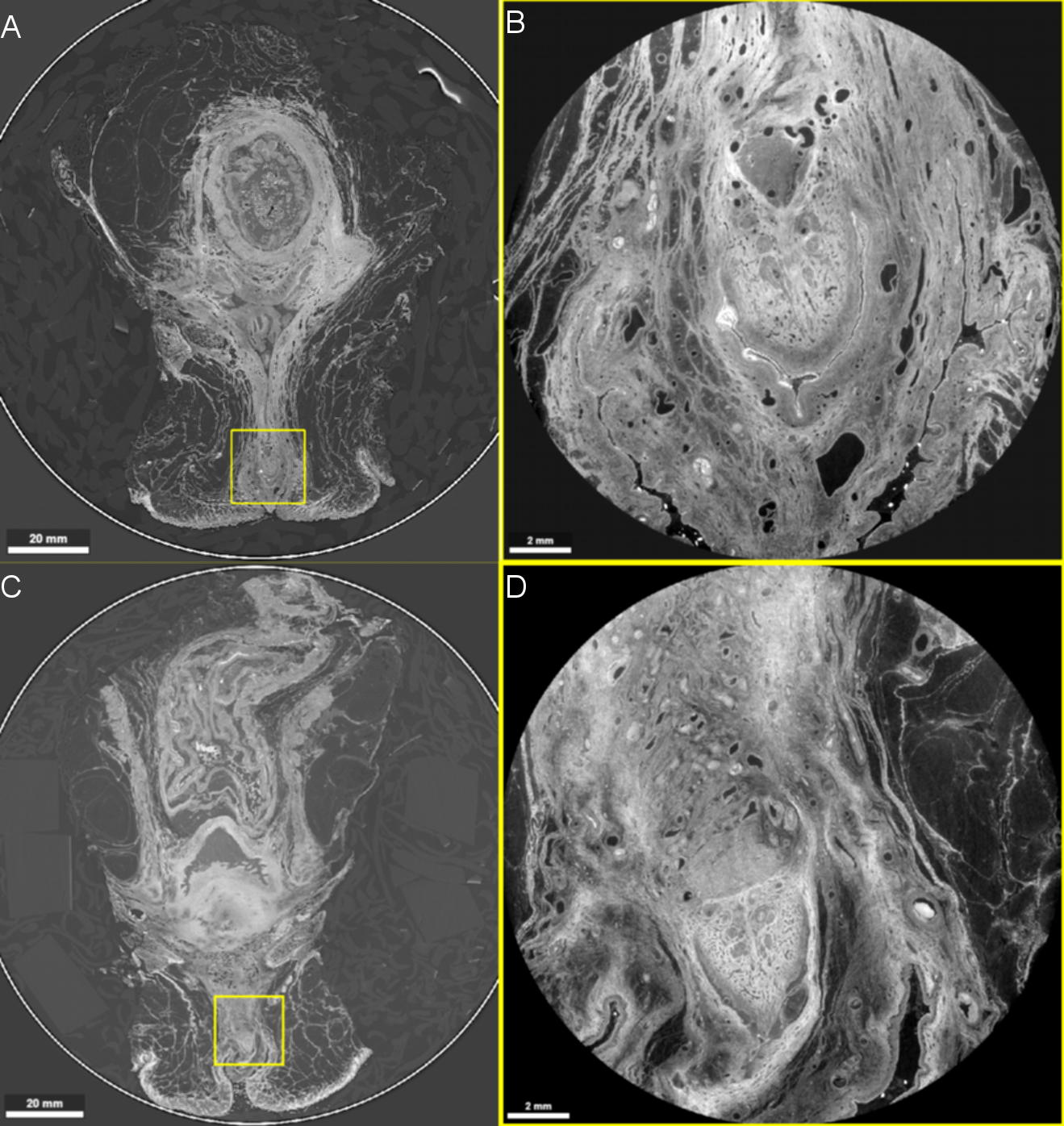

Figure S2. Synchrotron imaging at two voxel size settings. A. Field-of-view (FOV) for 20 micron setting (Subject 1. DOI: 10.15151/ESRF-DC-2379011888). B. FOV for 2 micron voxel size setting (Subject 1. DOI: 10.15151/ESRF-DC-2380206854). The position is noted with yellow rectangle in panel A. C. FOV for 20 micron setting (Subject 2. DOI: 10.15151/ESRF-DC-2380206870 ). D. FOV for 2 micron voxel size setting (Subject 2. DOI: 10.15151/ESRF-DC-2380206862). The position is noted with yellow rectangle in panel C. Scale bars are 20 m for panels A and C, 2 mm for panels B and D.

Supplementary Table 1

Size of the nerves within the clitoral glans

|  | Subject 1 | Subject 2 |
| --- | --- | --- |
| The diameter of maximum inscribed ball sorted from largest to smallest (µm) | 702.40 | 490.81 |
|  | 684.84 | 396.19 |
|  | 368.76 | 378.45 |
|  | 257.55 | 372.54 |
|  | 228.28 | 354.80 |
| Average (µm) | 448.365 | 398.56 |
| Standard deviation (µm) | 230.02 | 53.65 |

Supplementary Video 1

3D view of the clitoral glans innervation. The animation shows the dorsal nerve of the clitoris within the clitoral glans from Figure 2. Five large nerve trunks are shown within the glans, labelled in different colours. The surface of the clitoral glans is indicated in a grey line. The size of the bounding box is 5.7 mm x 7.9 mm x 7.3 mm.
