## Supplemental Table 2 for "Neuroanatomy of the clitoris"

| **Voxel size** | **20.06 µm** | **2.196 µm** |
|---|---|---|
| **Optic** | dzoom optic (Hasselblad 120 mm macro) 0.381x magnification | tandem 2X Nikkor 200 mm f/d 2 - Otus 100 mm f/d 1.4 |
| **Average energy** | 93 keV | 89 keV |
| **Filters** | sapphire 2.5 mm | sapphire 10 mm |
| | silver 0.1 mm | silver 0.2 mm |
| **Surface dose rate on** | 56.6 Gy/s | 8.6 Gy/s |
| **Integrated surface dose** | 11.5 kGy | 4.4 kGy |
| **Propagation distance** | 20 m | 2 m |
| **Sensor** | iris 15 (5056*1024) binning 2 by average | iris 15 (5056*2960) |
| **Single frame field of** | 50.71 * 10.27 $mm^2$ | 11.1 * 6.5 $mm^2$ |
| **Scintillator** | GAGG:Ce 1mm reflective | GAGG:Ce 50 µm reflective |
| **Source** | BM18 lateral beam 1.1 T (lateral shift by -16 mm at primary slits) | BM18 lateral beam 1.1 T (lateral shift by -10 mm at primary slits) |
| **Projection number** | 11000 | 15000 |
| **Jar mounting** | 140 mm diameter Jar immersed with | 140 mm diameter Jar immersed with |
| **Scan geometry** | quarter-acquisition helical scan with 5.5 mm vertical displacement per turn | half-acquisition, zseries with 4.5 mm vertical displacement between scans |
| **Reference geometry** | reference centered in the 2 mm thick polycarbonate tube filled with 70% ethanol, full integration of a complete rotation | reference off-axis in the 2 mm thick polycarbonate tube filled with 70% ethanol, partial integration with 30 angular portions along a single turn |
| **Scanning field of view** | 147 mm diameter, 137 mm height / 147 mm diameter, 164 mm height | 18.2 mm diameter, 24.5 mm height / 18.2 mm diameter, 47 mm height |
| **Exposure time** | 13 ms | 35 ms |
| **Number of scans / turns** | 2x24 helix turns + 2x2 flat scans / 2x29 helix turns + 2x2 flat scans | 5 scans / 9 scans |
| **Time per scan / turn** | 2.6 min per turn 2.3 h / 2.7 h | 9.6 min per scan 0.8 h / 1.6 h |
| **Reconstruction protocol** | processing with nightrail. ring corrections by partial average of projections over several turns, helical tomography reconstruction using filtered back-projection with single distance phase retrieval coupled with unsharp mask | processing with nightrail. ring corrections by partial average of projections over several turns, z-series tomography reconstruction using filtered back-projection with single distance phase retrieval coupled with unsharp mask |
| **Post-processing** | 16 bits conversion, exportation in jp2 format | 16 bits conversion, ring artefacts correction on reconstructed slices, exportation in jp2 format |
